# Circulating Th17.1 cells as candidate for prediction of therapeutic response to abatacept in patients with rheumatoid arthritis: exploratory research

**DOI:** 10.1101/593475

**Authors:** Shinji Maeda, Satoshi Osaga, Tomoyo Maeda, Norihisa Takeda, Shin-ya Tamechika, Taio Naniwa, Akio Niimi

## Abstract

T helper 17.1 cells (Th17.1) are highly pathogenic T cells in inflammatory diseases. This study aimed to identify Th cell biomarkers, including the analysis of Th17.1, that predict therapeutic response to abatacept in patients with rheumatoid arthritis. The circulating Th subsets among CD4+ T lymphocytes in 40 patients with rheumatoid arthritis before abatacept treatment were determined using multicolor flow cytometry. All patients received abatacept treatment for 24 weeks, and the change in disease activity score, including 28-joint count C-reactive protein, and the responsiveness of other indices to abatacept treatment were evaluated according the European League Against Rheumatism criteria [good responders, moderate responders, and non-responders]. The correlation between the abatacept responses and the Th subsets (baseline) was analyzed. Logistic regression analysis with inverse probability weighting method was conducted to calculate odds ratio adjusted for patient characteristics. The proportion of baseline Th17.1 cells was significantly lower in patients categorized as good responders than in those categorized as non-good responders (moderate responders and non-responders; p = 0.0064). The decrease in 28-joint count C-reactive protein after 24 weeks of abatacept therapy also showed a significant negative correlation with the proportion of Th17.1 cells. The adjusted odds ratio for achieving good response in patients with baseline Th17.1 level below the median value was 14.6 (95% confidence interval, 2.9–72.3; p = 0.0021) relative to that in the remaining patients. The proportion of Th17.1 cells at baseline is a good candidate for the prediction of response to abatacept treatment. These novel findings may represent an important step in the pursuit of precision medicine.

## Introduction

Advances in medicine and pharmaceutical technology have led to tremendous improvements in the treatment of rheumatoid arthritis (RA) [1]. Moreover, research in clinical human immunology has found new autoimmune cells. The development of potent anti-rheumatic drugs, particularly biological products, has helped in the improvement of clinical remission rates [2,3]. The targets of these biologics include both cytokines and T cells, which play key roles in the pathogenesis of RA. The therapeutic effect of abatacept (ABA), a strong inhibitor of T cells [4], has been shown to be equivalent to that of tumor necrosis factor α (TNF-α) inhibitor therapy [5–7].

The target lymphocytes of ABA, particularly CD4+ T cells, play a central role in the pathogenesis of RA, particularly in terms of acquired immunity, and in the induction of autoimmune response [4]. Research conducted on autoimmune mice models [8,9] has demonstrated the decisive role of Th17 in the pathogenesis of arthritis and autoimmune diseases. In humans, CCR6, which is a representative surface marker of Th17 [10], is a disease susceptibility gene of human RA [11]. Th17 is also involved in human RA pathology [12] by antigenically producing inflammatory cytokines, such as interleukin (IL)-17A, IL-17F, and IL-22. Unlike mice, these human Th17 cells have subpopulations. In particular, Th17.1 cells that are CD161+ CXC chemokine receptors (CXCR3)+ interferon-γ-producing Th17 cells, also referred to as classical Th1 or extinguish Th17 (ex-Th17) are a subgroup of Th17 cells found in humans [13] and are believed to be the most pathogenic among CCR6+ CD4+ T cells [12]. Th17 cells convert to inflammatory Th17.1 cells in an inflammatory milieu induced by cytokines, such as IL-1β, IL-23, TNF-α, and IL-12 [14]. Th17.1 cells have high expression levels of P-glycoprotein/multidrug resistance type 1 (MDR-1) and exhibit glucocorticoid resistance [13]. RA is characterized by joint destruction that is resistant to steroid treatment alone [15]. It is believed that the glucocorticoid resistance of Th17.1 cells is not attributable only to the function of MDR-1, and thus, further elucidation of the mechanism is expected in future. ABA is a strong inhibitor of these pathological T cells.

Although the therapeutic efficacy of ABA is well demonstrated, some patients are refractory to ABA treatment. The immune mechanisms that drive the chronicity of synovitis are multifactorial, such as adoptive immune pathways, innate immune pathways, stromal pathways, and systemic pathways [16]. Therefore, the contribution of immune factors other than T cells may influence the efficacy of ABA therapy. Among the various T cell subsets, some decrease in response to ABA treatment, whereas others do not [17]. ABA treatment has been reported to markedly reduce the proportion of T follicular helper cells and slightly decrease Th17 and activated regulatory T cells (Treg) cells. However, few studies have focused on the relationship between Th17.1 cells and therapeutic response to ABA in patients with RA. Moreover, there is a need to identify biomarkers that predict the therapeutic response. Identification of drug-specific biomarkers that predict therapeutic response is a desirable goal in the realm of personalized medicine [1]. Indeed, advances in cell analysis technology has raised prospects for the discovery of novel cellular immunological biomarkers that can predict treatment response in rheumatic diseases [18,19].

In the present study, we analyzed the proportion of each Th cell subset, including CXCR3+ Th17 cells (Th17.1), in the peripheral blood of patients with active RA before ABA treatment to explore early cellular biomarkers of response to ABA. We demonstrated the correlation between the proportion of Th17.1 cells and response to ABA therapy and proposed its potential use as a biomarker for predicting therapeutic response.

## Materials and Methods

### Ethics statement

This research was approved by the Ethics Review Committee of the Graduate School of Medicine, Nagoya City University. The study was conducted in compliance with the Declaration of Helsinki. Written informed consent was obtained from all patients.

### Participants

Patients with RA who fulfilled the classification criteria of the 1987 American College of Rheumatology rheumatoid arthritis classification (revised criteria of classification of RA) [20] and received ABA treatment at the Nagoya City University Hospital between 2009 to 2015 were eligible for inclusion. Inclusion criteria were as follows: 1) patients who consented to participate in this research, 2) patients who agreed to provide peripheral blood mononuclear cells (PBMCs) for immune-phenotyping analysis of Th subsets and Treg using multicolor flow cytometry, and 3) patients who did not achieve adequate improvement on previous treatment with at least one conventional synthetic disease-modifying anti-rheumatic drug (DMARDs).

Participants received intravenous ABA at 0 weeks, 2 weeks, 4 weeks, and every 4 weeks thereafter. The dose of ABA was based on body weight (BW) as follows: 500 mg for patients with a BW of <60 kg and 750 mg for those with a BW of 60 to 100 kg.

### Cell surface and intracellular staining and flow cytometry analysis

After obtaining informed consent, we obtained PBMCs of participants at baseline and at 4 and 24 weeks of ABA treatment. PBMCs were separated by density gradient centrifugation with Ficoll–Paque Plus (GE Healthcare, Uppsala, Sweden) and resuspended in flow cytometry buffer (Hank’s Balanced Salt Solution supplemented with 2% heat-inactivated fetal calf serum, 0.05% Sodium Azide, and 0.5% EDTA). Cells were stained for 30 min at 4°C under darkened conditions with the following fluorochrome labeled monoclonal antibodies: anti-CD4-AmCyan (Clone SK3, BD Biosciences, Franklin Lakes, Nj, USA), anti-CD25-APCCy7 (Clone M-A251, BD Biosciences), anti-CD45RA-FITC (Clone HI100, BD Biosciences), anti-Ki67-FITC (Clone B56, BD Biosciences), anti-CD196 (CCR6)-PE-Cyanine7 (Clone R6H1, eBioscience, San Diego, CA, USA), anti-CD161-PE (Clone HP-3G10, eBioscience), anti-CCR4-Alexa647 (Clone TG6/CCR4, eBioscience), anti-CD183 (CXCR3)-Pacific Blue (Clone G025H7, BioLegend), anti-forkhead box P3 (Foxp3)-PerCP-Cyamine5.5 (Clone PCH101, eBioscience), rat immune(Ig)G2a-PerCP-Cyamine5.5 antibody (eBioscience), and anti-CD45RO-APC (Clone UCHL1, TONBO Biosciences, San Diego, CA, USA). For the intracellular staining of Foxp3 and Ki67, the Foxp3-Staining Buffer Set (fixation/permeabilization and permeabilization buffers, eBioscience) was used according to the manufacturer’s protocol. Stained cells were washed twice using the flow cytometry buffer and resuspended for analysis using the Canto-II Flow cytometer (BD Bioscience) and the Diva software (BD Bioscience) and analyzed with FlowJo software (Tree Star). We defined the Th subset as follows:

Treg, CD4+ CD25+ Foxp3+; nonTreg, CD4+ Foxp3−; Th1, CXCR3+ CCR4− CCR6-nonTreg; Th2, CCR4+ CXCR3− CCR6− nonTreg; Th1&2, CXCR3+ CCR4+ CCR6− nonTreg; Th17, CCR6+ CD161+ CCR4+ CXCR3− nonTreg; and Th17.1, CCR6+ CD161+ CXCR3+ CCR4− nonTreg.

### Clinical assessment and evaluation of therapeutic response

Data pertaining to the following demographic and clinical variables were obtained from the medical records: age, sex, disease duration, use of corticosteroids, disease-modifying anti-rheumatic drugs, non-steroidal anti-inflammatory drugs, tender joint count, swollen joint count, patient global assessment [patient visual analog scale, 0–100 mm], physician global assessment [doctor’s visual analog scale, 0–100 mm], C-reactive protein (CRP) level, matrix metalloproteinase-3 (MMP-3) level, rheumatoid factor (RF) level, and anti-citrullinated protein/peptide antibody (ACPA) level.

Disease activity was assessed by calculating DAS28-CRP for each patient at each visit. The DAS28-CRP was calculated, and patients were categorized into the following four groups: remission, low (LDA), moderate (MDA), or high (HDA) disease activity according to the recommended formula (https://www.das-score.nl/das28/en/difference-between-the-das-and-das28/how-to-measure-the-das28/how-to-calculate-the-das28/alternative-validated-formulae.html). Because DAS28-CRP values are reportedly lower than those obtained in the original DAS28 assessment using erythrocyte sedimentation rate, a threshold of 4.1 (instead of the original 5.1) was used as the cut-off for HDA; 2.7 (instead of 3.2) as the cut-off for low disease LDA; and 2.3 (instead of 2.6) as the cut-off for remission [21]. The therapeutic response to ABA at 24 weeks was evaluated using the European League Against Arthritis (EULAR) response criteria (https://www.das-score.nl/das28/en/difference-between-the-das-and-das28/importance-of-das28-and-tight-control/eular-response-criteria.html), with 4.1 and 2.7 used as the thresholds for HDA and LDA, respectively. Briefly, patients were classified into three groups based on their 6-month DAS28-CRP and their absolute change from baseline according to the EULAR criteria as no response, moderate response, or good response. A good responder must demonstrate improvement of at least 1.2 units and achieve an absolute DAS28-CRP score of <2.7. A non-responder must demonstrate an improvement of ≤0.6 and have a final DAS28-CRP score of >4.1. Moderate responses fall between these data points. Furthermore, responsiveness to ABA treatment was evaluated using the following indicators: changes in disease activity before and after ABA treatment (ΔDAS28-CRP 0–24 weeks) and the evaluation of disease activity after 24 weeks of ABA treatment (remission, LDA, MDA, HDA).

After the initiation of ABA therapy, the clinical course was followed up for 24 weeks (every 4 weeks), and the correlation between responses to ABA treatment, RA disease activity, and the baseline proportion of Th subsets among CD4+ T lymphocytes (before treatment) was analyzed.

### MDR-1 activity assay

For the analysis of the MDR-1 activity of T cells, the fluorescent dye rhodamin 123 (Rh-123) was used according to the methods reported elsewhere [13]. Briefly, total CD4+ T cells were isolated using the Dynabeads CD4 positive T cell isolation kit (Invitrogen). Purified cells were 95%–98% pure as determined by flow cytometry analysis. Purified T cells in complete medium [DMEM (Gibco) supplemented with 10% FBS, 1% L-glutamine, 1% sodium pyruvate, 1% Hepes, and 1% Pen-Strep (all from Gibco) were loaded with Rh-123 (Sigma-Aldrich) at a final concentration of 1 μg/mL for 30 min on ice. Cells were then washed and moved to a 37°C incubator for 2 h. After an efflux period, cells were washed on ice in PBS, stained with surface markers (CD4, CD45RO, CXCR3, CCR6, and CD161), and washed again in PBS, and stained cells were kept on ice prior to flow cytometry analysis. Fluorescence reduction due to the emission of fluorescent dye by MDR-1 was confirmed by flow cytometry. For a negative control, 1 μM cyclosporine A (Sigma-Aldrich) was added to cells immediately before the incubation step.

### Statistical analysis

Mann-Whitney U test was used to assess between-group differences with respect to continuous variables, and Fisher’s exact test was used to assess between-group differences with respect to categorical variables. Kruskal–Wallis test was used for the analysis of differences in continuous variables between three groups. Friedman rank sum test and Wilcoxon signed rank test were used to analyze sequential changes in the proportion of each Th subset among CD4+ lymphocytes (0, 4, and 24 weeks) and Ki67 expression in each Th subset (0 and 4 weeks). Spearman rank correlation coefficient was used to assess correlation between two continuous variables. Stepwise variable selection method based on Akaike’s Information Criterion, Bayesian information criterion, and p-value was performed to identify the candidate Th subset that predicted ABA response.

The enrolled patients (n = 40) were divided into two groups based on the median proportion of Th17.1 cells among CD4+ T cells: Th17.1-lower (n = 20) and Th17.1-higher (n = 20).

To minimize the potential confounding effect due to baseline differences in patient characteristics between Th17.1-lower and higher groups, the inverse probability weighting (IPW) method, which is an application of the propensity score [22–24], was applied to compare the DAS28-CRP and good response rate of ABA treatment between the groups. Propensity scores for the IPW method were estimated using multivariate logistic regression analysis with the Th17.1 status (lower or higher) as the dependent variable and using the following baseline characteristics as independent variables: age, sex, DAS28-CRP (baseline), RF, ACPA RA disease duration (years), history of biological DMARDs, prescription of methotrexate (MTX), and glucocorticoids. The discriminative power of the propensity score was quantified by the c statistic corresponding to the area under the receiver operating characteristic curve. Next, each patient background variable was compared under the correction by the IPW method using the weighted Mann–Whitney test, the weighted *t*-test, and the weighted chi-squared test, and the respective p-values were calculated. The effect of Th17.1 on patient ABA response was evaluated using estimated odds ratios (OR) and 95% confidence intervals (CIs). All calculated p-values were two-sided, and p-values < 0.05 were considered statistically significant for all analyses. Statistical analyses were performed with the R-software version 3.3.3 (R Development Core Team, Vienna, Austria) and EZR version 1.35 (Saitama Medical Center, Jichi Medical University, Saitama, Japan) [25], which is a graphical user interface for R (The R Foundation for Statistical Computing, Vienna, Austria). The following R-software packages were used for statistical processing and creation of graphs and tables: survey (version 3.31-5) [26], aod (version 1.3), weights (version 0.85), ggplot2 [27], and corrplot (version 0.77).

## Results

### Baseline characteristics of patients

Table 1 shows the baseline demographics and clinical characteristics of the enrolled patients (N = 40). The disease activity of RA was high in the study population (median DAS28-CRP, 4.43; simplified disease activity index, 23.8). ACPA-positive patients accounted for 60% of the study population, and they were relatively older (median age, 70.5 years). With respect to use of concomitant drugs, 77.5% of the patients were taking MTX, whereas 60% were taking glucocorticoids. With respect to medication history, only 32.5% of the patients had a history of treatment with biological DMARDs.

**Table 1.** Clinical characteristics of patients at baseline.

|  |  |
| --- | --- |
| Patient characteristics | Overall n = 40 |
|  | Median [IQR], or, (%) |
| Age, year old | 70.5 [60.8, 74.6] |
| Sex: male/female, (%) | 8/32 (20.0/80.0) |
| Disease duration (years) | 4.2 [1.5, 15.9] |
| DAS28-CRP | 4.43 [4.02, 5.01] |
| SDAI score | 23.8 [20.3, 28.8] |
| CRP, mg/dl | 1.02 [0.48, 2.27] |
| MMP-3 (ng/ml) | 167.7 [100.2, 290.2] |
| ACPA (negative/positive, (%)) | 16/24 (40.0/60.0) |
| negative, (%) | 16 (40.0) |
| Low positive, (%) | 5 (12.5) |
| High positive, (%) | 19 (47.5) |
| RF (negative/positive, (%)) | 14/26 (35.0/65.0) |
| negative, (%) | 14 (35.0) |
| Low positive, (%) | 9 (22.5) |
| High positive, (%) | 17 (42.5) |
| Concomitant methotrexate, n (%) | 31 (77.5) |
| MTX mg ( mean (sd)) | 9.4 (2.9) |
| Concomitant glucocorticoid, n (%) | 24 (60.0) |
| prednisolone, mg (mean, (sd)) | 5.8 (3.6) |
| Concomitant tacrolimus, n (%) | 3 ( 7.5) |
| Concomitant NSAIDs, n (%) | 20 (50.0) |
| Biologic DMARDs naïve, n (%) | 27 (67.5) |
| Pulmonary complications associated with RA, n (%) | 18 (45.0) |

This table shows patient baseline demographics. Data are presented as median [IQR, interquartile range], mean [SD], or frequency [%].

DAS28-CRP, disease activity score 28-joint count C-reactive protein; SDAI, simplified disease activity index; CRP, C-reactive protein; NSAIDs, non-steroidal anti-inflammatory drugs; MMP-3, matrix metallo-proteinase 3; ACPA, anti-citrullinated protein antibody; RF, rheumatoid factor; MTX, methotrexate; DMARDs, disease modified anti-rheumatic-drug; Low positive, less than 3 times normal upper limit among positive; High positive, more than 3 times the normal upper limit.

Between-group differences with respect to median values determined using the Mann– Whitney U test, whereas those with respect to percentage values were determined using the Fisher’s exact test.

### Characterization of Th17.1 in patients with RA

We analyzed the subtype of peripheral blood T cells before and four weeks after ABA treatment (Fig 1a); furthermore, in each cell group, Ki67 expression was determined by flow cytometry as a cell proliferation marker.

**Fig 1.**
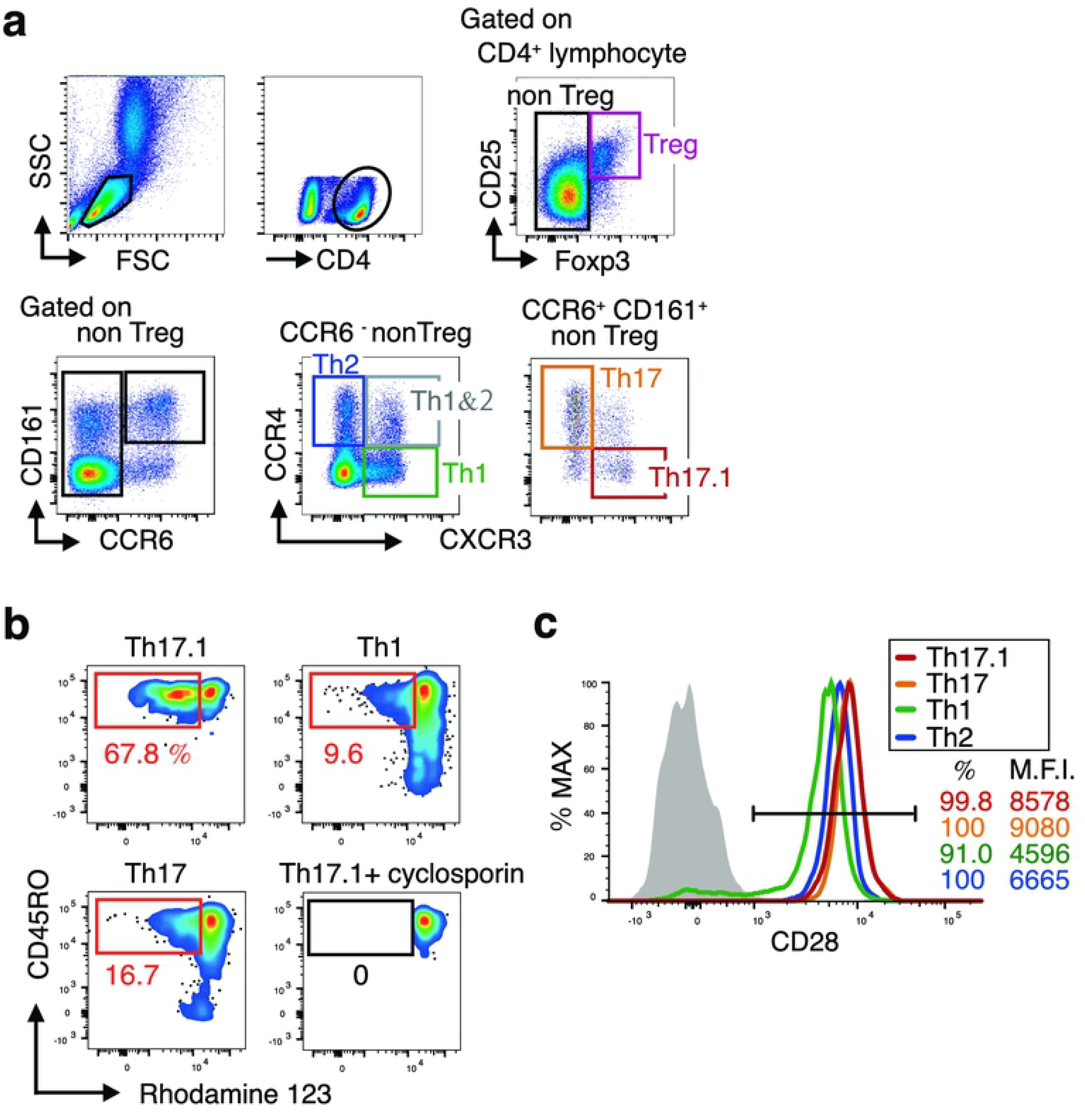
Characteristic of circulating Th17.1 cells in patients with rheumatoid arthritis (RA). **a**. Flow cytometry plots explaining the gating strategy for the identification of peripheral blood Treg, Th1, Th2, Th1&2, Th17, and Th17.1 subpopulations. CD4+ T cell subsets in the peripheral blood of adult patients with RA were analyzed using flow cytometry. b. The MDR1 activity of indicated Th subset assessed using multicolor flow cytometry with rhodamine 123 (Rh-123). Total CD4+ T cells isolated from peripheral blood were labeled with Rh123. After a 1-h efflux period at 37°C in the presence of vehicle (Dimethyl sulfoxide) or MDR1 inhibitors (cyclosporine A), cells were stained with antibodies to CCR6, CXCR3, and CD45RO, and Rh-123 efflux of each Th subsets was analyzed using flow cytometry. Data shown are flow cytometry plots representing three independent experiments performed on cells isolated from different donors with RA. c. Proportion of cells with CD28 expression within the indicated CD4+ T cell subpopulations of patients with RA. Data are representative of at least three independent experiments.

Th17.1 was the smallest subset of CD4+ cells (median 1.17%; interquartile range 0.71–1.93) (S1 Fig). Next, the expression of MDR-1, one of the major features of Th17.1, was confirmed using Rh-123 (Fig 1b). The results showed that MDR-1 was highly expressed only in Th17.1 cells and not in the Th1 and Th17 cells, as reported so far. The expression rate of CD28, an inhibitory target in ABA treatment, in the Th17.1 cells was as high as that in the others (>99%) (Fig 1c).

### Difference between early changes in proliferation status of Th subsets

Changes in the proportions of Th subset among CD4+ T cells before ABA treatment and 4 weeks after treatment were confirmed to evaluate the effect of ABA treatment on each Th subset. However, noticeable changes were not observed (Fig 2a). Therefore, we next analyzed the expression of Ki67 in the cells to confirm the early effects of ABA on each Th cell subset (Fig 2b). Before ABA treatment, the proportion of Ki67 positive cells among each Th subset was different; particularly, the expression rate in Th17.1 cells was remarkably lower than that in the other subsets (S2 Fig, Figs 2b and 2c). In contrast, the expression rate of Ki67 in Treg cells was relatively higher, which suggests that Treg is active during cell proliferation in patients with RA. Next, the Ki67 positivity rate for each Th subset was determined after ABA treatment for 4 four weeks and compared with the baseline. In only 4 weeks, the proportion of Ki67 positive cells was significantly reduced in all subsets other than the Th17.1 cells (Fig 2c). The change in Ki67 expression in Th17.1 cells was not statistically significant (p = 0.39).

**Fig 2.**
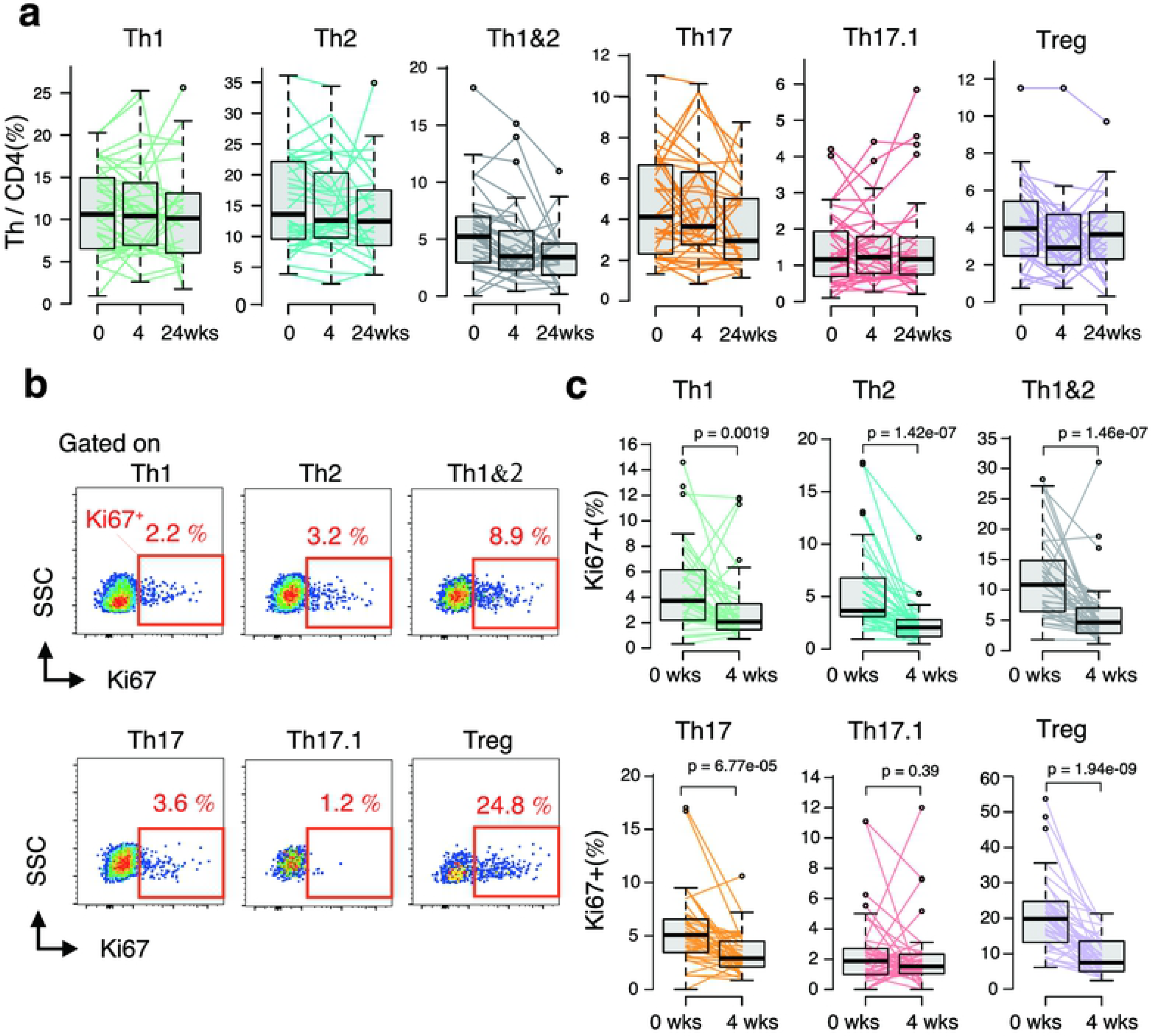
Early change in cell proliferation state of each Th subset by ABA treatment. Peripheral blood mononuclear cells of patients with RA were obtained at baseline and at 4 and 24 weeks of abatacept treatment (0 and 4 weeks, n = 40; 24 weeks, n = 29). a. Sequential changes in the proportion of each T cell subset among CD4 T cells in peripheral blood induced by ABA treatment. Data were analyzed using Friedman rank sum test. b. Flow cytometry plot showing the frequency of Ki67 expression in the indicated CD4+ Th cells in patients with RA assessed using intracellular staining of Ki67 antigen and analyzed using multicolor flow cytometry. c. Sequential changes in the proportion of Ki67 expression in each Th subsets induced by ABA treatment (0 and 4 weeks, n = 40). Data were analyzed using Wilcoxon signed rank test.

### Therapeutic response to ABA and baseline Th17.1

ABA treatment was continued for 24 weeks, and the progress of disease activity in each patient and the responsiveness to ABA treatment were evaluated. Subsequently, we analyzed the correlation between ABA response and the Th subset at baseline. A remarkable finding was that the proportion of baseline Th17.1 cells among CD4+ T cells in good responders was significantly lower than that in poor responders (p = 0.0064) (Fig 3a). In contrast, no significant difference was observed with respect to the other Th subsets.

**Fig 3.**
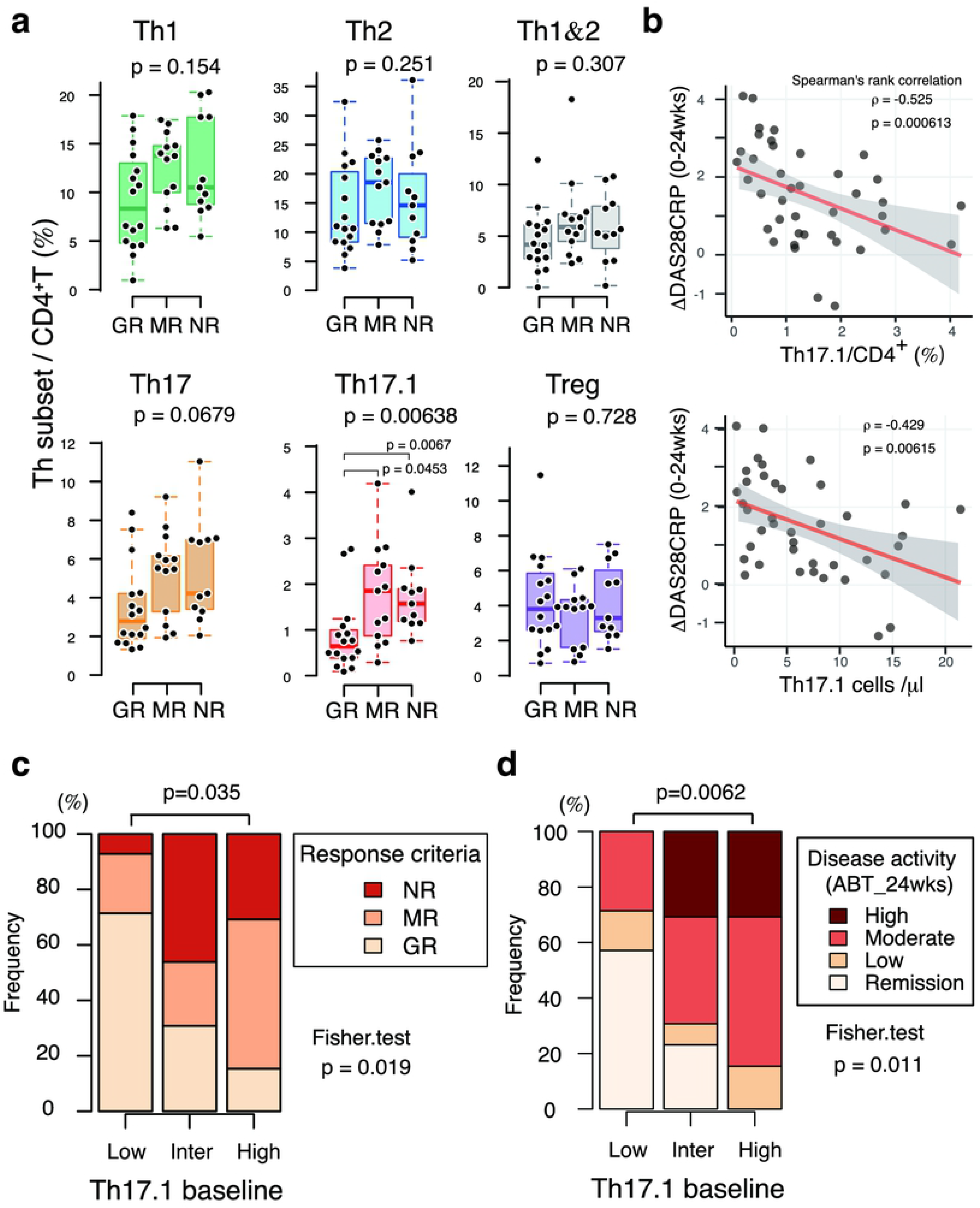
Clinical significance of Th17.1 levels in abatacept treatment response. ABA treatment was continued for 24 weeks. Subsequently, correlation between ABA response and the Th subset at baseline was analyzed. The following indicators were used to evaluate response to ABA treatment: changes in DAS28-CRP scores from baseline at 24 weeks after ABA treatment (ΔDAS28-CRP 24 weeks), disease activity evaluation after 24-week ABA treatment [remission and low (LDA), moderate (MDA), and high (HDA) disease activity], and treatment response evaluation using the EULAR response criteria [good responder (GR), moderate responder (MR), and non-responder (NR)]. a. ABA treatment response after 24 weeks was evaluated as GR (n = 14), MR (n = 13), and NR (n = 13). The proportion of indicated Th subsets among CD4 + lymphocytes at baseline in each group was plotted and displayed as box plot. b. Scatter plot shows the ratio or absolute number of Th17.1 cells at baseline and ΔDAS28-CRP 24 weeks. Regression line (red line) and 95 CI of the regression line (gray zone) are also shown in the plot. c. d. Patients were stratified into three groups (low, intermediate, and high) based on tertiles of the proportion of Th17.1. Hundred percentage stacked bar chart shows the EULAR response rate (c) and the disease activity score (d) at 24 weeks after the initiation of ABA treatment in three groups. Data were analyzed using the Kruskal–Wallis and Mann–Whitney tests for between-groups comparisons with Bonferroni correction (a), Spearman’s rank correlation coefficient (b), and Fisher’s exact test and Bonferroni correction for paired comparisons (c, d).

The attenuation of disease activity (ΔDAS28-CRP) after 24 weeks of ABA treatment also showed a significant negative correlation with Th17.1 (both percentage and absolute number) (Fig 3b).

To assess the clinical relevance of the correlation between Th17.1 cells and ABA response, we divided the patients into two groups (lower and higher) using the median Th 17.1 proportion (/CD4+) as cut-off (Table 2), and into three groups (lowest, intermediate, and highest) based on tertiles. ABA treatment response in each group was analyzed. Treatment response was significantly different in the three groups (p = 0.019) (Fig 3c). The treatment response in the Th17.1-lower (lowest) group was significantly better than that in the higher (highest) group (S3a–S3b Figs). On analysis of the trend of disease activity at 24 weeks, a lower proportion of Th17.1 cells at baseline was associated with a higher percentage of remission (Fig 3d). Remission rates in the Th17.1-lower and lowest groups were 55% and 57.1%, respectively, and no remission was observed in the higher and highest groups.

**Table 2.** Clinical characteristics of the Th17.1-lower and Th17.1-higher patients at baseline

| Patient characteristics | Th17.1 / CD4+ T cells |  | p |
| --- | --- | --- | --- |
|  | Lower<br>n = 20 | Higher<br>n = 20 |  |
| Age, year old | 70.5 [60.8, 74.2] | 71.0 [63.4, 74.7] | 1.00 |
| Sex: male/female, (%) | 6/14 (30.0/70.0) | 2/18 (10.0/90.0) | 0.24 |
| Disease duration (years) | 3.4 [0.9, 6.4] | 6.7 [2.5, 19.1] | 0.07 |
| DAS28-CRP | 4.50 [3.77, 4.91] | 4.43 [4.22, 5.02] | 0.71 |
| SDAI score | 22.8 [18.5, 28.7] | 24.3 [21.6, 29.2] | 0.31 |
| CRP, mg/dl | 1.18 [0.57, 2.52] | 0.82 [0.48, 2.16] | 0.81 |
| MMP-3 (ng/ml) | 206.3 [104.6, 316.1] | 152.2 [94.8, 267.8] | 0.50 |
| ACPA (negative/positive, (%)) | 9/11 (45.0/55.0) | 7/13 (35.0/65.0) | 0.75 |
| negative, (%) | 9 (45.0) | 7 (35.0) |  |
| Low positive, (%) | 3 (15.0) | 2 (10.0) | 0.67 |
| High positive, (%) | 8 (40.0) | 11 (55.0) |  |
| RF (negative/positive, (%)) | 7/13 (35.0/65.0) | 7/13 (35.0/65.0) | 1.00 |
| negative, (%) | 7 (35.0) | 7 (35.0) |  |
| Low positive, (%) | 5 (25.0) | 4 (20.0) | 1.00 |
| High positive, (%) | 8 (40.0) | 9 (45.0) |  |
| Concomitant methotrexate, n (%) | 17 (85.0) | 14 (70.0) | 0.45 |
| MTX mg ( mean (sd)) | 10.2 (2.5) | 8.3 (3.1) | 0.07 |
| Concomitant glucocorticoid, n (%) | 13 (65.0) | 11 (55.0) | 0.75 |
| prednisolone, mg (mean, (sd)) | 5.5 (4.3) | 6.1 (3.0) | 0.72 |
| Concomitant tacrolimus, n (%) | 0 (0.0) | 3 (15.0) | 0.23 |
| Concomitant NSAIDs, n (%) | 10 (50.0) | 10 (50.0) | 1.00 |
| Biologic DMARDs naïve, n (%) | 14 (70.0) | 13 (65.0) | 1.00 |
| Pulmonary complications associated with RA, n (%) | 7 (35.0) | 11 (55.0) | 0.34 |

Enrolled patients (n = 40) were stratified into two groups based on the median proportion of Th17.1 cells among CD4+ T cells: Th17.1-lower (n = 20) and Th17.1-higher (n = 20). The table shows clinical features and differences of patient subgroups of Th17.1-lower and Th17.1-higher at baseline. Data presented as median [IQR, interquartile range] or mean [SD], or frequency [%].

DAS28-CRP, disease activity score 28-joint count C-reactive protein; SDAI, simplified disease activity index; CRP, C-reactive protein; NSAIDs, non-steroidal anti-inflammatory drugs; MMP-3, matrix metallo- proteinase 3; ACPA, anti-citrullinated protein antibody; RF, rheumatoid factor; MTX, methotrexate; DMARDs, disease modified anti-rheumatic-drug; Low positive, less than 3 times normal upper limit among positive; High positive, more than 3 times the normal upper limit.

Between-group differences with respect to median values determined using Mann–Whitney U test, and those with respect to percentage values determined using Fisher’s exact test.

We assessed whether changes in objective biomarkers of arthritis [serum CRP, MMP-3)] after ABA treatment for 24 weeks were different between the Th17.1 lower and higher groups. A significant reduction in serum CRP (S4a Fig) and MMP-3 levels (the significant high-titer class) was observed in the Th17.1-lower group (S4b and S4c Figs).

Subsequently, we performed receiver operating characteristic curve analysis to determine the optimal threshold level of Th17.1 associated with good response or remission at 24 weeks (S5 Fig). A cut-off level of 1.09% (Th17.1 cells/CD4+ cells) was associated with 79.2% sensitivity and 81.2% specificity for GR and 75.9% sensitivity and 100% specificity for remission.

### Th17.1 and patient background factors

Differences in the clinical features between the Th17.1-lower and Th17.1-higher groups at baseline were evaluated; however, no significant differences were observed. Next, because CD4+ T cells play an important role in the pathogenesis of RA, we assessed the correlation between various patient characteristics, disease activity, Th subset, and ABA therapeutic response using Spearman’s rank correlation coefficient (S6 Fig). The baseline disease activity (DAS28-CRP baseline) showed a strong correlation with serum CRP, ACPA, RF, and MMP-3 levels. In the Th subset, although baseline disease activity showed a negative correlation with Treg, no significant correlation was observed with other Th subsets. The disease duration of RA showed a positive correlation between Th1&2, Th17, and Th17.1 cells. In the Th subset analysis, Th17.1 showed a strong correlation with Th1 and Th17. Analysis of the correlation between patient characteristics and ABA response revealed a strong correlation of ΔDAS28-CRP (0–24 weeks) with baseline DAS28-CRP and age. However, none of the patient characteristics showed a significant correlation with EULAR response criteria and disease activity after 24 weeks (S6 Fig, S1 Table). Therefore, in this study, we found no meaningful association between the background characteristics of patients and therapeutic response to ABA. However, baseline levels of Th17.1 and Th17 subsets showed a significant association with all three indices of ABA response, and baseline Th1 level was significantly associated only with disease activity after 24 weeks. Among these three Th subsets, the Th17.1 subset showed the most significant association with ABA response.

Among the patient background factors and Th subsets, we selected and narrowed down the candidate variables to construct an optimal model for prognostic prediction using stepwise variable selection in multivariate analysis (S2 Table). In all multivariate analyses, only Th17.1 was selected and showed the most significant association after adjustment for potential confounders. Given the limited number of cases for adjusting confounding factors by multivariate analysis, we used the IPW method to reduce the number of confounders and to analyze the adjusted effect of baseline Th17.1 on ABA therapeutic response. The c statistic, the discriminative power of propensity score (PS) for Th17.1-lower group was 0.735 (95% CI 0.576–0.894). With IPW, all patient background factors that are shown in S3 Table were more evenly adjusted between Th17.1-lower and higher, including RA disease duration, which was not significantly different but tended to correlate. Of note, even after adjustment for different covariate distributions for both groups, there was a significant difference between Th17.1-lower and higher groups with respect to ΔDAS28-CRP (0–24 weeks) and disease activity after 24 weeks (DAS28-CRP 24 weeks) (Fig 4a). The effect of Th17.1-lower on ABA good response as compared to that of Th17.1-higher was evaluated by estimated OR with 95% CIs, after adjustment by IPW. In this study, good responders and patients with low disease activity after 24 weeks were equivalent. As a result, in the Th17.1-lower group, OR for achieving good response was 14.6 (95% CI, 2.9–72.3; p = 0.0021) (Fig 4b, S4 Table). The proportion of Th17.1 cells among CD4+ T cells at baseline was a good predictor of ABA treatment response.

**Fig 4.**
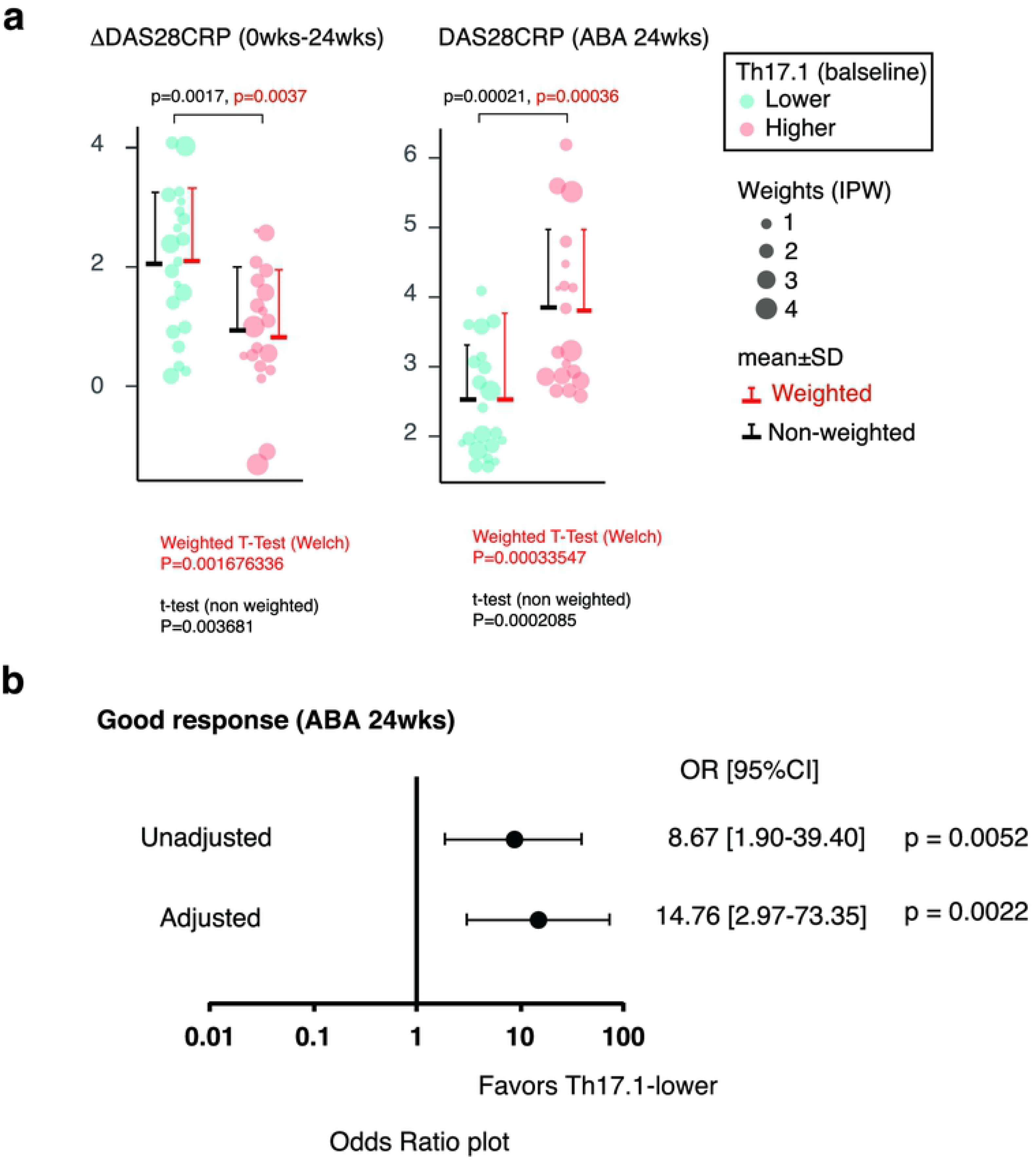
Prediction of therapeutic response to ABA based on the proportion of Th 17.1 at baseline. **a**. The difference in ABA therapeutic response between Th17.1-lower (binary by median) and Th17.1-higher after adjustment of patient background factors using inverse probability weighting (IPW). The size of the balloon plot indicates the weighting using the IPW method in each case. The red lines indicate the weighted mean (horizontal line) and SD (vertical line) after IPW adjustment. Black lines indicate non-weighted mean (horizontal line) and SD (vertical line). Data were analyzed using the weighted (red) or non-weighted (black) *t*-test. b. The adjusted odds ratio for achievement of GR with baseline Th17.1-lower relative to Th17.1-higher. Logistic regression analysis using the IPW method to calculate the odds ratio adjusted for patient characteristics. Forest plot shows unadjusted and adjusted odds ratios and 95% CI and p-value. ABA, abatacept; IPW, inverse probability weighting; SD, standard deviation; OR, odds ratio; CI, confidence interval

## Discussion

In the present study, we found that baseline Th17.1 levels may be a prognostic predictor of ABA treatment in patients with RA. Th17 also correlated with ABA therapeutic response; however, Th17.1 showed a stronger correlation with ABA therapeutic response. Among Th cells that cause antigen-specific responses, the identification of cells that correlate with ABA therapeutic response is the most novel finding of this study.

A key novelty aspect of this study was the inclusion of analysis of cell surface CD161, intracellular forkhead box P3 (Foxp3), and Ki 67 in flow cytometry analysis. Although CD161 is not included as a surface marker in the international standard human immune-phenotyping method [28], it is particularly important as a surface marker of pathogenic Th17 and Th17.1 subsets. In addition, CCR6 and CCR4 are also expressed in Treg; therefore, intracellular Foxp3 was stained to precisely exclude regulatory T cells. Analysis of the expression of Ki67 allowed us to capture the early change in cell proliferation status of each Th subset induced by ABA treatment. When compared with the reported standard method, there are certain limitations in the interpretation of this research method and results; however, it is important to analyze Th17.1 more precisely.

In this study, Th17.1 showed a significant correlation with the patient background characteristics such as disease duration. Even after adjustment for patient background characteristics using the IPW method, the baseline proportion of Th17.1 subset significantly predicted ABA responsiveness. For the identification of cellular immunological biomarkers that predict therapeutic response by flow cytometric analysis, adjustment for patient background characteristics is important because target cells themselves may interact with patient background factors other than therapeutic effect. However, because of the limited number of samples owing to the handling of living cells and the associated cost burden, it is difficult to adjust for confounding factors using multivariate analysis. In such settings, the use of the IPW method to adjust for patient background characteristics is a well-accepted practice.

Previous studies have found only few factors that adequately predict the therapeutic response to ABA in RA treatment. These include CRP [29], presence or absence of MTX combination, history of use of other biologics [30], positivity for ACPA and RF [31], and age [32]. In this study, no significant correlation was observed between these known baseline patient characteristics and the proportion of peripheral blood Th17.1 cells. None of the known patient characteristics was found to predict ABA treatment response. Furthermore, the following lymphocyte-related indices, which are independent of Th17.1, were reported as prognostic factors: proportion of terminally differentiated effector memory cells among CD8 T cells [33] and the proportion of CD28 negative T cells [34,35]. However, these lymphocyte subsets were not analyzed in this study. Therefore, the correlation between these known lymphocytes and Th17.1 was not clarified, which is a study limitation.

Our data also showed interesting results pertaining to intracellular Ki67 expression in each Th subset. The expression level of Ki67 in Th17.1 cells at baseline was significantly lower than that in the other subsets. Moreover, the positivity rate of Ki67 in Th subsets, except for Th17.1, decreased markedly in the early stage of ABA treatment. These findings suggest that Th17.1 is less susceptible to ABA with respect to cell proliferation. Considering that the mechanism of action of several immunosuppressive drugs (such as DMARDs) involves the inhibition of lymphocyte proliferation, immunosuppressive drugs are less likely to target Th17.1 because Th17.1 is non-proliferating in active patients with RA undergoing conventional synthetic therapy with DMARDs.

In contrast to Th17.1 cells, the difference in Treg levels did not predict the response to ABA treatment despite the difference in Treg proportions in the patients at baseline. Immunological mechanisms of RA pathology, such as the disruption of autoimmune tolerance, onset and persistence of inflammation, and joint destruction, are extremely complex. Even in the contribution of T cells to the persistence of inflammation, the importance of the involvement of dysfunction in the suppression (regulatory) side, such as Treg, and the existence of inflammatory T cells that can resist Treg suppression is also conceivable. Treg expresses cytotoxic T-lymphocyte antigen-4 (CTLA-4) [36], which is an immunosuppressive functional molecule common to both Treg and ABA. It was inferred that the inactivation of the immune system by CTLA-4-Ig strongly supplemented the difference in the amount of Treg and the influence of endogenous Treg for the arthritis condition will be reduced. In contrast, when inflammatory cells that are resistant to suppression by these CTLA-4 molecules are present at baseline, these levels are likely to affect disease activity after the ABA treatment.

These results pose a simple question of whether Th17.1 has a causal relationship with treatment resistance. In previous reports, Th17.1 cells were described as the most pathologic inflammatory cells among the CCR6 positive T cells [12]. Both Th17 and Th17.1 cells are present in the synovial fluid of patients with RA; however, Th17.1 cells are particularly more abundant in the synovial fluid than in the peripheral blood [37]. Compared with Th1 and Th17, Th17.1 cells produce more types of pro-inflammatory cytokines, such as IL-17A, granulocyte macrophage colony-stimulating factor, interferon-γ, and TNF-α, which are associated with rheumatoid inflammatory conditions. More specifically, treatment with the neutralizing antibody of the inflammatory cytokine granulocyte macrophage colony-stimulating factor, which is strongly produced by Th17.1, is effective in RA [38]. Therefore, we envisage a certain role of Th17.1 in the activity of RA even during ABA treatment in non-responders. CTLA-4 molecule is a strong negative regulator of T cell immune response [39,40] and plays the central role in Treg’s regulatory function [36]. More interestingly, it was recently reported that ex-Th17, which has the same phenotype as Th17.1, is not restricted by Treg suppression [41]. Based on these facts and the results of this study, it is suggested that Th17.1 cells play a role in disrupting immune tolerance by CTLA-4 and Treg. Currently, it is possible to induce clinical remission in some patients with RA using biological DMARDs or targeted synthetic DMARDs. However, when targeting immunologic remission, immunological factors as therapeutic targets still remain unknown. Immunological remission can be achieved by the inactivation or suppression of inflammation-sustaining pathological factors, such as autoreactive-T-cells, via endogenous immune regulatory mechanisms, such as CTLA-4 or Treg. Although further biological studies are needed to assess whether Th17.1 is resistant to the repression of CTLA-4-Ig, our study suggests that Th17.1 cells serve as potential novel therapeutic targets for achieving immunologic remission.

## Conclusions

The present study demonstrated that the proportion of circulating Th17.1 cells showed differences in immunological quality that determine the therapeutic response to ABA in patients with RA. A vast array of anti-rheumatic drugs is currently available for the treatment of RA. The identification of the most appropriate drug for individual patients is a key imperative to achieve early improvement. The identification of Th17.1 as a good candidate biomarker of the therapeutic response to ABA may represent an important step in the pursuit of precision medicine.

## Author contributions

SM conceived, designed the study, performed data analysis, and wrote the manuscript. SO contributed to determining statistical methods for data analysis and statistical programming and interpretation and assisted in the preparation of the manuscript. SM and TM performed all flow cytometric analyses, and SM and TT conducted an experiment of MDR-1 activity assay. ST helped with obtaining patient’s blood samples. All other authors have contributed to data collection and interpretation and have critically reviewed the manuscript. All authors have approved the final version of the manuscript.

## Acknowledgements

We thank Mrs. Y. Sato and R. Uemura for the help in collecting patients’ blood samples. The authors would like to thank the patients who participated in this study and all members of our laboratory for the meaningful discussions.

## Supporting information

**S1 Fig. The proportion of cells with Ki67 expression among circulating CD4+ Th subsets in patients with RA**PBMCs from RA (n = 26, before abatacept treatment) were stained for CD4, CXCR3, CCR4, CD161, CCR6, CD25, Foxp3, and Ki67 mAbs and analyzed using flow cytometry. The percentage of Ki67+ cells in the indicated Th subsets are shown by box plot. Data were analyzed using Kruskal–Wallis test followed by the Mann– Whitney U test using Bonferroni correction.

**S2 Fig. Change in cell proliferation state of Th17 and Th17.1 subsets by ABA treatment.** The graph shows changes in the proportion of cells with Ki67 expression among each Th subset induced by ABA treatment at various time-points (0, 4, and 24 weeks, n = 29).

Data were analyzed using Friedman rank sum test. Wilcoxon signed rank test with Bonferroni correction was used for post-hoc paired comparisons.

**S3 Fig. Th17.1 level and successive changes in disease activity score.**

**a**. The line graph shows the transition of the disease activity (DAS28-CRP) of RA in the Th 17.1-lower and Th 17.1-higher groups before and after ABA treatment (at 4, 12, and 24 weeks). P-values (vs. Th17.1-higher) were determined with Mann–Whitney U test using Bonferroni correction. **b**. 100% stacked bar chart shows successive changes in DAS28-CRP in the Th17.1-lower, Th17.1-intermediate, and Th17.1-high groups before and after ABA treatment (at 4, 12, and 24 weeks).

ABA, abatacept; DAS28-CRP, disease activity score 28-joint count C-reactive protein; REM, remission; LDA, low disease activity; MDA, moderate disease activity; HDA, high disease activity.

**S4 Fig. Th17.1 level and successive changes in CRP and MMP-3 levels.**

**a. b.** The line graphs show the transition of serum C-reactive protein (CRP) and metalloproteinase-3 (MMP-3) of rheumatoid arthritis in Th 17.1-lower and Th 17.1-higher groups before and after ABA treatment (at 4, 12, and 24 weeks). P-values (Th17.1-lower vs. Th17.1-higher) were determined using the Mann–Whitney U test.

**c**. 100% stacked bar chart shows MMP-3 titer (normal, moderate, and high) in Th17.1-low and Th17.1-high groups after ABA treatment at 24 weeks. P-values (Th17.1-lower vs. Th17.1-higher) were determined using Fisher’s exact test.

ABA, abatacept; CRP, C-reactive protein; MMP-3, metalloproteinase-3; Normal, within normal limit; Moderate titer, less than 3 times normal upper limit; High titer, more than 3 times normal upper limit.

**S5 Fig. Estimation of Th17.1 cut-off value at baseline to predict ABA therapeutic response using ROC curve**.

**a**. ROC curve showing a cut-off Th17.1 (% in CD4+) level of 1.1% discriminated between GR and non-GR (MR or NR) at 24 weeks, with 79.2% sensitivity and 81.2% specificity.

**b**. ROC curve showing a cut-off Th17.1 level of 1.1% discriminated between REM and non-REM at 24 weeks, with 75.9% sensitivity and 100% specificity.

ROC, receiver operating characteristic; AUC, area under the curve; GR, good response; MR, moderate response; NR, no response; REM, remission.

**S6 Fig.** Correlation coefficient matrix plot shows the correlation (Spearman’s correlation coefficient, ρ) of patient background factors, indicated T cell subset at baseline, and ABA therapeutic response indicators with significance levels (p-value).

**S1 Table. Differences in baseline clinical characteristics between EULAR-GR and non-GR patients**.

**S2 Table. Exploratory analysis for optimal Th subset as the predictor of ABA treatment response using multivariate analysis**.

**S3 Table. Adjusted patient characteristics of Th17.1-lower and Th17.1-higher patients by IPW**.

**S4 Table. Logistic regression analysis using the IPW method to calculate odds ratio adjusted for patient characteristics**.

